# Scalable mass-spectrometry-based molecular phylogeny with TreeMS2

**DOI:** 10.64898/2026.02.27.708420

**Authors:** Michel Dierckx, Charlotte Adams, Julia M. Gauglitz, Wout Bittremieux

## Abstract

Molecular phylogeny is a well-established method for inferring evolutionary relationships from DNA and RNA sequences. Here, we extend this concept beyond genetic information by applying phylogeny-like analysis to proteomic and metabolomic mass spectrometry data, capturing relationships based on the realized molecular phenotype. The resulting phenotype-derived trees can be directly compared with conventional genetic-based trees to identify where molecular phenotypes reflect evolutionary history and where they diverge due to functional adaptation, regulation, or environmental influence.

To enable this analysis, we introduce TreeMS2, a computational tool that constructs similarity matrices by directly comparing tandem mass spectrometry (MS/MS) spectra between samples. By bypassing spectrum annotation, TreeMS2 enables rapid, unbiased comparisons. Across diverse datasets, TreeMS2 reconstructs biologically meaningful relationships. In proteomics, phenotype-derived trees recapitulate established taxonomy, with deviations pinpointing sample handling errors. In single-cell proteomics our method distinguishes cell types despite sparse and noisy measurements and in metabolomics it resolves major biochemical divisions and fine-scale compositional structure. Together, these results establish TreeMS2 as a scalable, annotation-independent framework for deriving molecular relationships from raw MS/MS data.

## Introduction

Molecular phylogeny has traditionally relied on DNA and RNA sequences [1], which provide a highly informative record of evolutionary history based on the genetic blueprint. In contrast, the functional biochemical state of an organism as captured by the proteome and metabolome represents the realized molecular phenotype, integrating the influences of genetic variation, epigenetic regulation, and environmental context. Mass spectrometry (MS)-based molecular phenotyping can therefore reveal patterns of biochemical convergence and divergence that deviate from purely sequence-based relationships, offering new insight into adaptation, niche specialization, and convergent evolution.

Advances in mass spectrometry have generated vast public repositories of proteomics (e.g., PRIDE [2], MassIVE, ProteomeXchange [3]) and metabolomics (e.g., GNPS/MassIVE [4], MetaboLights [5], Metabolomics WorkBench [6]) data, containing billions of spectra from thousands of species, tissues, and environments. Despite coverage that in many cases rivals or even exceeds genomic databases, these resources remain largely untapped for phylogenetic analysis, particularly in taxa where genomic data are sparse or incomplete. Large-scale integration of proteomics and metabolomics has the potential to yield unified molecular phylogenies, due to shared biology and biochemical processes, spanning bacteria, archaea, and eukaryotes, bridging organismal domains in ways that are often inaccessible to sequence-based approaches. However, unlike sequence-based phylogenetics, which benefits from decades of methodological development, there is no widely adopted computational framework for reconstructing phylogeny directly from MS data at scale.

Existing methods such as compareMS2 for proteomics and Qemistree for metabolomics demonstrate the feasibility of deriving relationships from molecular profiles but remain limited in scope. CompareMS2 [7,8] compares mass spectra to compute similarity between MS runs and has been applied in small-scale studies for species identification in foods [9,10] and quality control [11], yet its lack of scalability limits its applicability to large datasets. Qemistree [12], by contrast, operates at the feature level, converting untargeted metabolomics data into molecular fingerprints via *in silico* annotation via SIRIUS [13] and CSI:FingerID [14] before building hierarchical feature trees. While powerful, its reliance on accurate fingerprint predictions makes it vulnerable to the low-quality or unannotated spectra common in large public datasets, and it does not produce sample- or taxon-level phylogenies suitable for cross-study comparisons. As a result, current approaches are constrained to localized comparisons and cannot efficiently operate on thousands of MS runs needed for large-scale biochemical phylogeny.

Here we introduce TreeMS2, a scalable bioinformatics framework for phylogenetic analysis from MS-based molecular data. TreeMS2 can process heterogeneous proteomics and metabolomics data, computes large-scale similarity matrices suitable for phylogenetic reconstruction and dimensionality reduction, and scales to hundreds to thousands of MS runs comprising millions of spectra, enabling analyses that were previously computationally prohibitive. Using a unified computational backbone, it supports both proteomic and metabolomic data, enabling multi-modality applications. We demonstrate the capabilities of TreeMS2 through diverse use cases, and provide it as open-source software at https://github.com/bittremieuxlab/TreeMS2, establishing a practical foundation for integrating molecular and evolutionary landscapes.

## Results

### Scalability of TreeMS2 for large-scale molecular phylogeny

We developed TreeMS2 as a scalable bioinformatics tool for constructing molecular phylogenies directly from large-scale MS data (**Figure 1**). The method represents each tandem mass (MS/MS) spectrum as a binned vector, applies sparse random projections for dimensionality reduction, and uses approximate nearest-neighbour search to rapidly identify similar spectra across datasets. Sample-to-sample similarity is defined as the average fraction of spectra in each sample that have at least one close match in the other sample, then these values are converted into a symmetric distance matrix suitable for downstream analyses. This approach enables the direct transformation of raw spectral data into phylogenetic trees or low-dimensional embeddings without requiring molecular identification.

**Figure 1.**
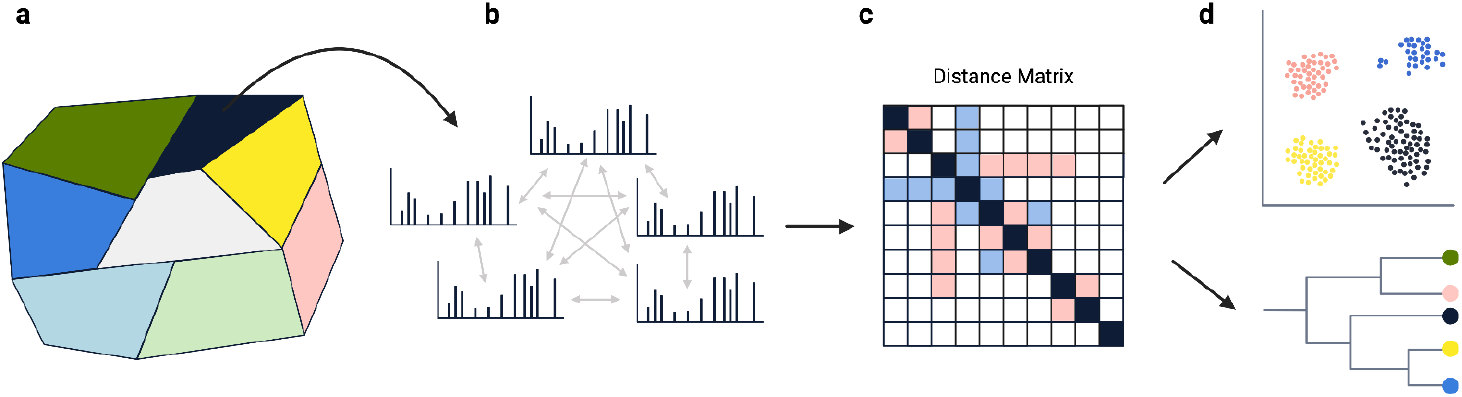
Overview of the TreeMS2 workflow. Raw MS/MS spectra from proteomics or metabolomics datasets are converted into binned vector representations and projected into a lower-dimensional space using sparse random projections. (**a**) These vectors are indexed for efficient approximate nearest-neighbour search, enabling (**b**) rapid spectrum-to-spectrum comparisons across large datasets. Pairwise sample similarities are computed as the average fraction of spectra in each sample with at least one close match in the other. (**c**) The resulting symmetric distance matrix can be used for (**d**) multiple downstream analyses, including phylogenetic tree reconstruction and dimensionality reduction methods such as UMAP for visualizing molecular relationships.

While the underlying similarity concept is inspired by compareMS2 [7,8], TreeMS2 introduces key algorithmic advances that enable analyses at scales unattainable by existing methods. In particular, spectrum vectorization combined with approximate nearest-neighbour search [15,16] reduces the computational complexity from a quadratic dependence on the number of spectra to a near-linear scaling in practice. This allows TreeMS2 to process datasets containing millions to hundreds of millions of MS/MS spectra, whereas compareMS2 is limited to small datasets due to its requirement for exhaustive pairwise spectral comparisons (**Figure 2**).

**Figure 2.**
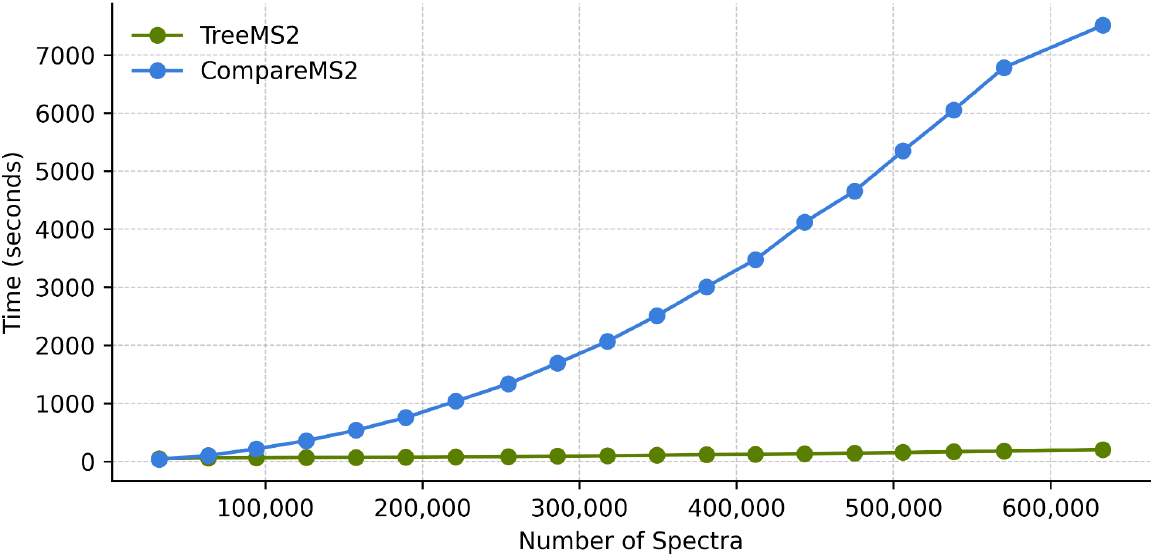
Runtime comparison of TreeMS2 and compareMS2 across increasing dataset sizes. Total processing time is shown as a function of the number of input MS/MS spectra. TreeMS2 exhibits near-linear scaling, maintaining practical runtimes even for large datasets, whereas compareMS2 scales quadratically and quickly becomes computationally intractable. Benchmarks were performed on identical hardware using subsets of the bacterial proteomics dataset [17].

Because TreeMS2 operates directly on raw tandem mass spectra, it is inherently multi-modal: the same workflow applies to proteomics and metabolomics datasets without domain-specific modifications. The method does not rely on peptide, protein, or metabolite identification, and is therefore unaffected by incomplete reference databases, which is a major advantage for environmental samples, under-studied taxa, and untargeted metabolomics datasets where identification rates are often low [18].

The output distance matrix generated by TreeMS2 is compatible with a wide range of analytical frameworks. Beyond phylogeny reconstruction, it can be used in ordination analyses such as principal component analysis (PCA) and uniform manifold approximation and projection (UMAP) [19], as well as ecological statistics such as permutational multivariate analysis of variance (PERMANOVA). This flexibility enables both evolutionary and ecological interpretations from the same underlying computation, bridging functional and phylogenetic perspectives.

### Bacterial proteomics phylogeny recovers taxonomy and flags data anomalies

We applied TreeMS2 to a bacterial proteomics dataset comprising 303 proteomes from 119 genera and five phyla [17]. The 38SPD dataset contains over 13 million MS/MS spectra and was processed in under 3 and a half hours and was thus easily tractable for TreeMS2 (**Supplementary Table 1**). By contrast, compareMS2 could not complete the analysis at all due to its quadratic scaling, confirming the practical advantages of TreeMS2 for datasets containing hundreds of proteomes and millions of spectra.

The resulting molecular phylogeny showed strong agreement with established bacterial taxonomy: the overall topology closely matched recognized phylum-level groupings and also recovered relationships at the class, order, and genus level (**Figure 3**). Quantitatively, proteomic distances were strongly correlated with taxonomic relatedness (Mantel ρ = 0.665, p < 9.999e-05). Notably, this agreement was achieved without genome sequence information or reference protein databases, as TreeMS2 operated entirely on the raw spectral content.

**Figure 3.**
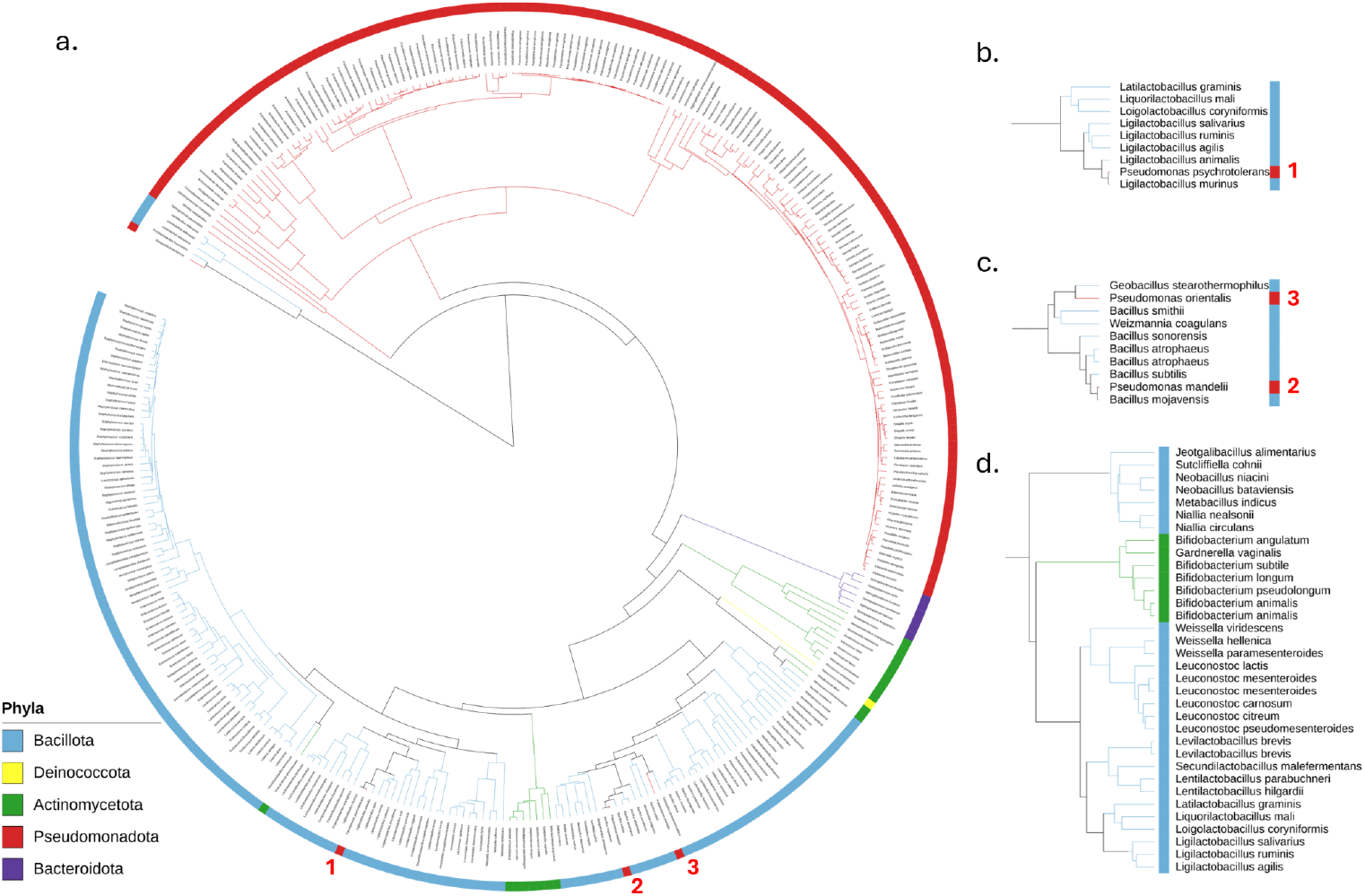
TreeMS2-derived phylogeny for 303 bacterial proteomes. (**a**) The tree was constructed from over 13 million MS/MS spectra spanning 119 genera and five phyla, using raw spectral similarity without genome sequence or protein database information. Major phyla are indicated by coloured branches. The topology shows strong agreement with established bacterial taxonomy, recovering relationships from the phylum to the genus level. Three *Pseudomonas* species (*P. psychrotolerans* (1), *P. mandelii* (2), and *P. orientalis* (3)) are highlighted and shown in more detail in **b** and **c**; all are unexpectedly distant from their close relatives due to sample-handling errors. (**d**) A highlighted section of the tree displaying the clustering of Bifidobacteria Bacillota instead of the other Actinomycetota.

Although most relationships were taxonomically consistent, a few species deviated from their expected positions. A notable case involves *Pseudomonas psychrotolerans, Pseudomonas orientalis*, and *Pseudomonas mandelii*, which appeared unexpectedly distant from their close relatives (**Figure 3**). For *P. psychrotolerans* and *P. orientalis*, the original study reported that the samples were mistakenly sourced from the wrong wells [17], providing a clear explanation. While no such annotation was made for *P. mandelii*, its displacement in the TreeMS2 phylogeny suggests a similar sample-handling error, which may have gone unnoticed when relying solely on peptide and protein identifications but becomes apparent from the raw MS/MS data. This indicates that TreeMS2 can naturally detect sampling errors and MS runs with low spectral quality, providing a built-in mechanism for quality control in large-scale analyses.

When searching *P. psychrotolerans* and *P. orientalis* against the protein sequence database of their closest neighbour in the phylogenetic tree (**Figure 3b-c**) we observed a drastic increase in the annotation rates, respectively from 0.5% and 0.1% to 21% and 18%. This supports the hypothesis that samples were sourced from the wrong wells. In addition, *P. mandelii*, which was not reported to be sourced from the wrong well, was similarly displaced in the phylogenetic tree. When searching *P. mandelii* against the fasta of its closest neighbour in the tree, again a drastic increase in the annotation rate was observed from 0.9% to 21%. TreeMS2 is thus not only able to detect these kinds of errors, it can help hypothesize what occurred, such as determining which species was likely sampled instead.

Another noticeable discrepancy in the tree is that a group of Actinomycetota are clustered between the Bacillota rather than with other Actinomycetota. This separately clustered group contains all 5 Bifidobacteria and *Gardnerella vaginalis*, also known as *Bifidobacterium vaginalis* (**Figure 3d**). One possible explanation is that Bifidobacteria are more functionally similar to Bacillota rather than the other Actinomycetota. To investigate this, we searched the species in the BacDive database and looked into the isolation sources [20]. The 5 Bifidobacteria were all found in feces or in sewage and *Gardnerella vaginalis* was found in urine and vaginal secretion. In contrast, none of the other Actinomycetota were found in feces, urine or any other isolation sources related to the gastrointestinal or urogenital tract. However, within the Bacillota we were able to find 33 species in feces, sewage or the gastrointestinal tract, including some well known gut microbes like *Enterococcus faecium* and *Enterococcus faecalis*. In addition, 8 Bacillota species were found in urine or the urogenital tract, 17 Bacillota species were found in blood, and 18 Bacillota species were found in food. Of the 24 remaining Bacillota species, 9 were not found in BacDive, and 15 were found in plants, soil, or animals. The fact that the isolation sources of Bacillota are more similar to Bifidobacteria compared to other Actinomycetota indicates that Bifidobacteria might have a more similar function resulting in more similar proteomes.

TreeMS2 was also used to analyze 46 bacterial isolates from dairy products for which ground truth bacterial species were obtained with 16S rRNA sequencing [17]. This dataset contained over 800 thousand MS/MS spectra and, combined with the 38SPD dataset, was processed in under 5 hours (**Supplementary Table 1**). These 46 of the food isolates have a genus that is present in the 38SPD dataset analyzed above and thus can be found in the phylogeny tree produced by TreeMS2 (**Figure 3**) and for 32 of these even the bacterial species are present. We trained a nearest neighbor classifier to predict the genus and for all 46 food isolates the genus was correctly predicted. The correct species was predicted for 31 out of 32 food isolates. The only misidentification was *Raoultella planticola* which should have been *Raoultella ornithinolytica*. The distance of the food isolate was the smallest to *Raoultella planticola* with 3.11, the second closest neighbor was *Raoultella ornithinolytica* with a distance of 3.32. On average, the difference between the distance to the first and second neighbor is 6.04, indicating that this was a more difficult distinction to make as the two species are highly similar.

### TreeMS2 scales from bacterial to complex proteomes

To evaluate TreeMS2 on larger and more complex proteomes, we analyzed data from a “kingdom of life” proteomics study [21], containing a total of 114 species: 14 viruses, 19 archaea, 48 bacteria, and 33 eukaryotes. The original study performed single runs for all prokaryotes and used offline fractionation generating eight LC–MS/MS fractions per species for eukaryotes because their proteomes are substantially larger and more complex. TreeMS2 includes native functionality to assign multiple input files to the same biological group, allowing the eight fractions for each species to be easily combined during analysis. Unfortunately, not all files could be mapped to their corresponding species and some raw files could not be converted to MGF (**Supplementary Table 2**). Consequently, only 79 species were covered: 11 viruses, 3 archaea, 32 bacteria, and 33 eukaryotes.

Despite the increased complexity, with millions of MS/MS spectra per species, TreeMS2 processed the data efficiently, requiring less than 13 hours to process the dataset containing over 56 million MS/MS spectra (**Supplementary Table 1**). This demonstrates that the same computational pipeline can handle datasets ranging from small bacterial proteomes to large eukaryotic proteomes without algorithmic modification or performance bottlenecks. The phylogenetic tree generated by TreeMS2 closely matches the established evolutionary relationships among the 79 species (**Figure 4**; Mantel ρ = 0.611, p < 9.999e-05). These results indicate that TreeMS2 can robustly recover known phylogenetic structure from highly complex proteomes while maintaining scalability.

**Figure 4.**
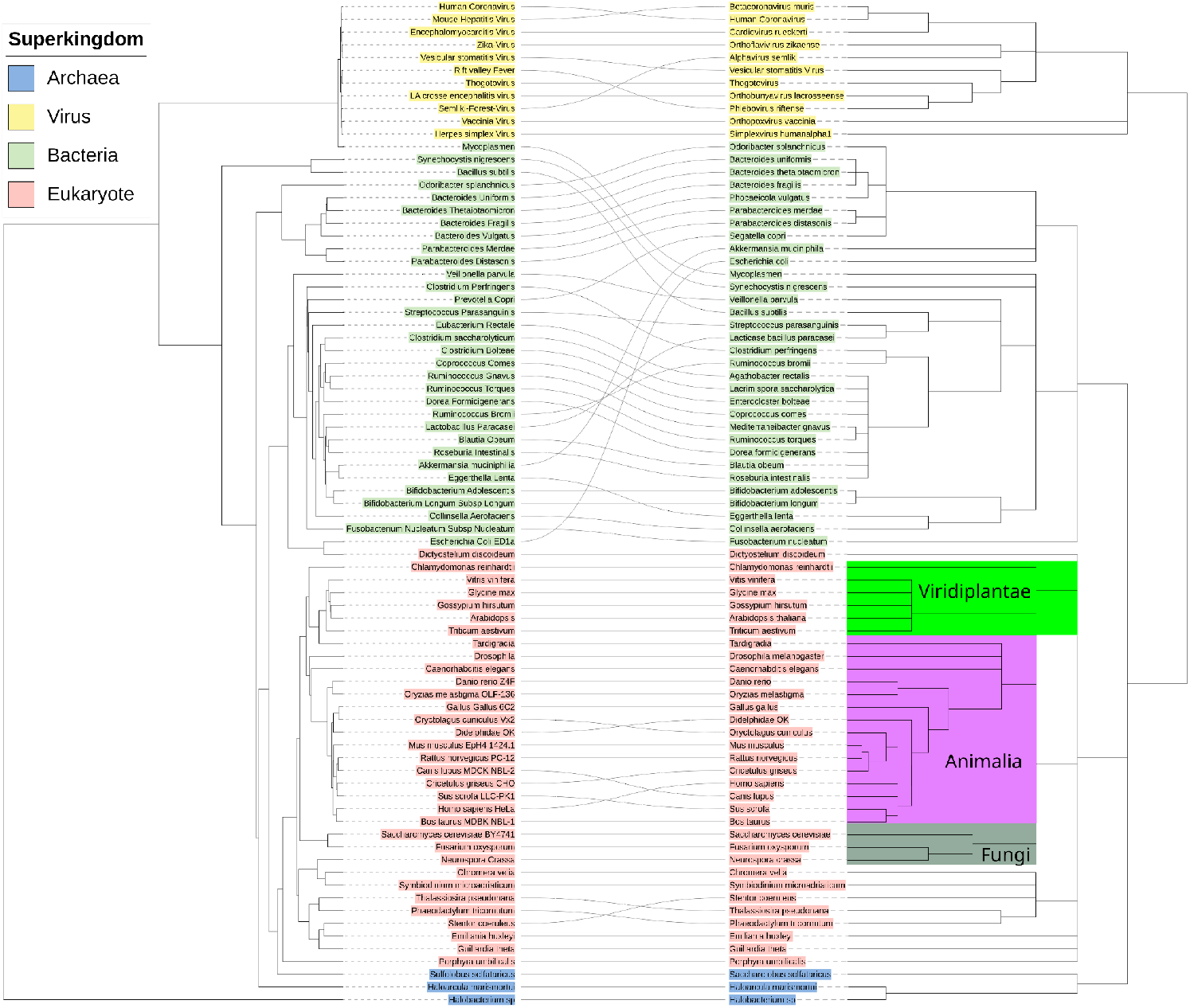
Mirror tanglegram of the TreeMS2 derived phylogeny compared to NCBI taxonomy tree. The TreeMS2 tree (left) was constructed from over 56 million MS/MS spectra, using raw spectral similarity without genome sequence or protein database information. The branch lengths of the NCBI taxonomy tree (right) are based on the taxonomic ranks. Superkingdom and kingdoms (specifically Viridiplantae, Animalia, and Fungi) are highlighted. The topology shows strong agreement with established taxonomy.

The TreeMS2 derived phylogeny closely matches the established evolutionary relationships, with the 46% of first neighbours matching between the two trees (**Supplementary Table 3**). In addition, the species are clustered according to larger groups. For example, viruses, bacteria, and eukaryotes are in distinct clusters. Within the eukaryotes the plants, animals, and fungi are also clustering together, as shown in **Figure 4**.

There are some interesting discrepancies to highlight. For example, *E. coli* and *Dictyostelium discoideum*, commonly referred to as social amoeba or slime mold, are clustered together even though they are evolutionary quite distinct. Our hypothesis is that *E. coli*, and thereby *E. coli* peptides, were present in the *D. discoideum* sample, as it was the food source on the agar plate they were growing on. To test this hypothesis, we searched the *D. discoideum* samples against a protein sequence database containing both *E. coli* and *D. discoideum*, expecting a relatively large fraction of the spectra to be annotated as *E. coli* peptides. The resulting annotation rate of 28% at 1% FDR with 80% of the annotated spectra originating from *D. discoideum* peptides and 20% from *E. coli* peptides corroborates our hypothesis.

Another interesting discrepancy is that even though mycoplasma is a bacterium it is closer to viruses rather than bacteria. One explanation could be that they are functionally quite similar to viruses, as they are small, lack a cell wall, and have the smallest genome for self-replicating organisms. For years after their discovery, they were even thought to be viruses, because they passed through the usual bacterial filters [22].

## Single-cell proteomics reveals differentiation trajectories

We next evaluated TreeMS2 on single-cell proteomics (SCP) data generated using the Chip-Tip workflow, which was designed for deep SCP analysis by minimizing sample losses during preparation and employing narrow-window data-independent acquisition (DIA) on an Orbitrap Astral mass spectrometer [23]. The dataset comprised human induced pluripotent stem cells (hiPSCs) and 53 embryoid body (EB) cells profiled during undirected differentiation, with up to 4,700 proteins quantified in hiPSCs and 6,200 in large EB cells. The dataset contains over 20 million spectra and was processed in under four hours (**Supplementary Table 1**).

TreeMS2 analysis revealed a clear separation between hiPSCs and EB cells (**Figure 5**). Consistent with the original study, some EB cells were as distant from each other as they were from hiPSCs, reflecting substantial heterogeneity within the EB population. Mapping the six clusters identified in the original study [23] onto the TreeMS2 embedding showed strong correspondence, indicating that the original differentiation patterns are reproducible directly from raw spectral similarity.

**Figure 5.**
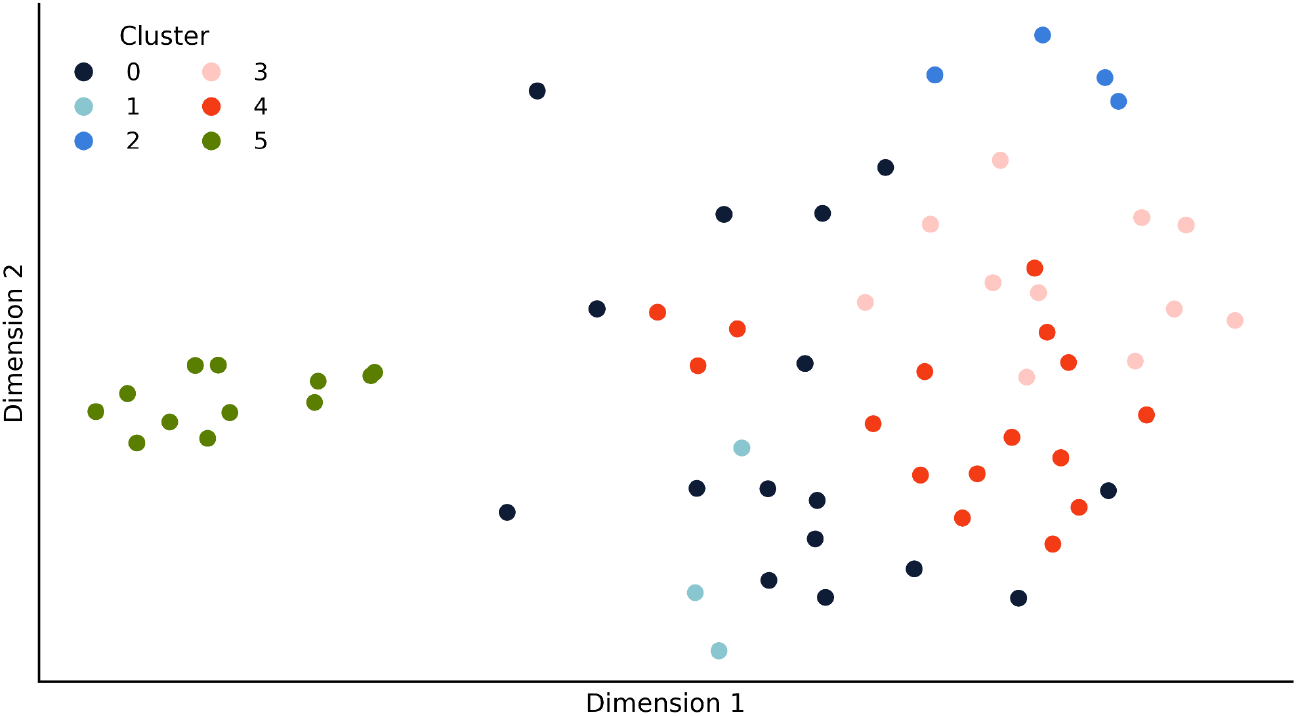
MDS embedding of SCP data based on TreeMS2 spectral similarity. Each point represents one cell, positioned according to a multidimensional scaling projection of the TreeMS2 distance matrix. Cells are colored by the six clusters reported in the original study [23]. The projection shows a clear separation between hiPSCs (cluster 5; green) and EBs (clusters 0-4), while also recapitulating the within-EB structure, with some EB cells as distant from each other as from hiPSCs.

Two aspects of this analysis are particularly noteworthy. First, it demonstrates that TreeMS2 can be applied to DIA data, whereas the previous use cases, as well as alternative tools for mass spectrometry-based similarity, have been limited to data-dependent acquisition (DDA). Second, it shows that the method is effective even for highly complex and potentially noisy data such as single-cell proteomics, which is a medically important class of data. SCP data present unique challenges, including low and variable signal intensity, a high proportion of missing peptide identifications, and increased stochasticity in detection [24]. Nevertheless, TreeMS2 was able to sensitively detect biologically meaningful structure without relying on derived peptide or protein identifications and quantifications, instead leveraging the complete raw MS/MS spectral content. This is particularly advantageous when protein quantification tables are sparse, noisy, or incomplete; conditions that are especially common in SCP experiments.

### Molecular diversity of the global food metabolome

Finally, as the input data for treeMS2 are MS/MS spectra in MGF format, a common mass spectrometry-based raw data format, we applied TreeMS2 to a large untargeted metabolomics dataset. The dataset comprises over 3,500 food samples from the Global FoodOmics project, spanning a wide range of plant- and animal-derived products [25], and containing over 4 million spectra. Each food item was profiled by untargeted tandem mass spectrometry, and the dataset includes rich metadata categorizing foods within a hierarchical ontology. This dataset differs substantially from the previous examples, both in molecular composition and in analytical modality, and was processed in less than two hours (**Supplementary Table 1**) with minor adjustments to the configurations.

Analysis of raw MS/MS spectra, without requiring metabolite annotation, is particularly advantageous in untargeted metabolomics, where annotation rates are often low [18] and discarding unannotated features can lead to substantial information loss. Furthermore, it is especially relevant in natural product research, where molecular diversity is vast—far exceeding that of, for example, the human metabolome—and reference libraries capture only a small fraction of existing molecules [26]. By working at the spectral level, TreeMS2 bypasses these coverage limitations and incorporates the complete molecular signal into the analysis, maximizing data utilization.

TreeMS2 analysis revealed molecular diversity patterns across multiple biological and functional scales. The UMAP embedding (**Figure 6**) generated from the TreeMS2 distance matrix provides a low-dimensional representation of dietary molecular diversity, in which local spatial proximity tends to reflect similarity in biochemical composition between foods. This representation allows intuitive navigation of complex metabolome data, enabling, for example, the identification of compositional gradients between food categories or the detection of unusual items whose molecular signatures deviate from their expected group. Such a representation provides an exploratory overview.

**Figure 6.**
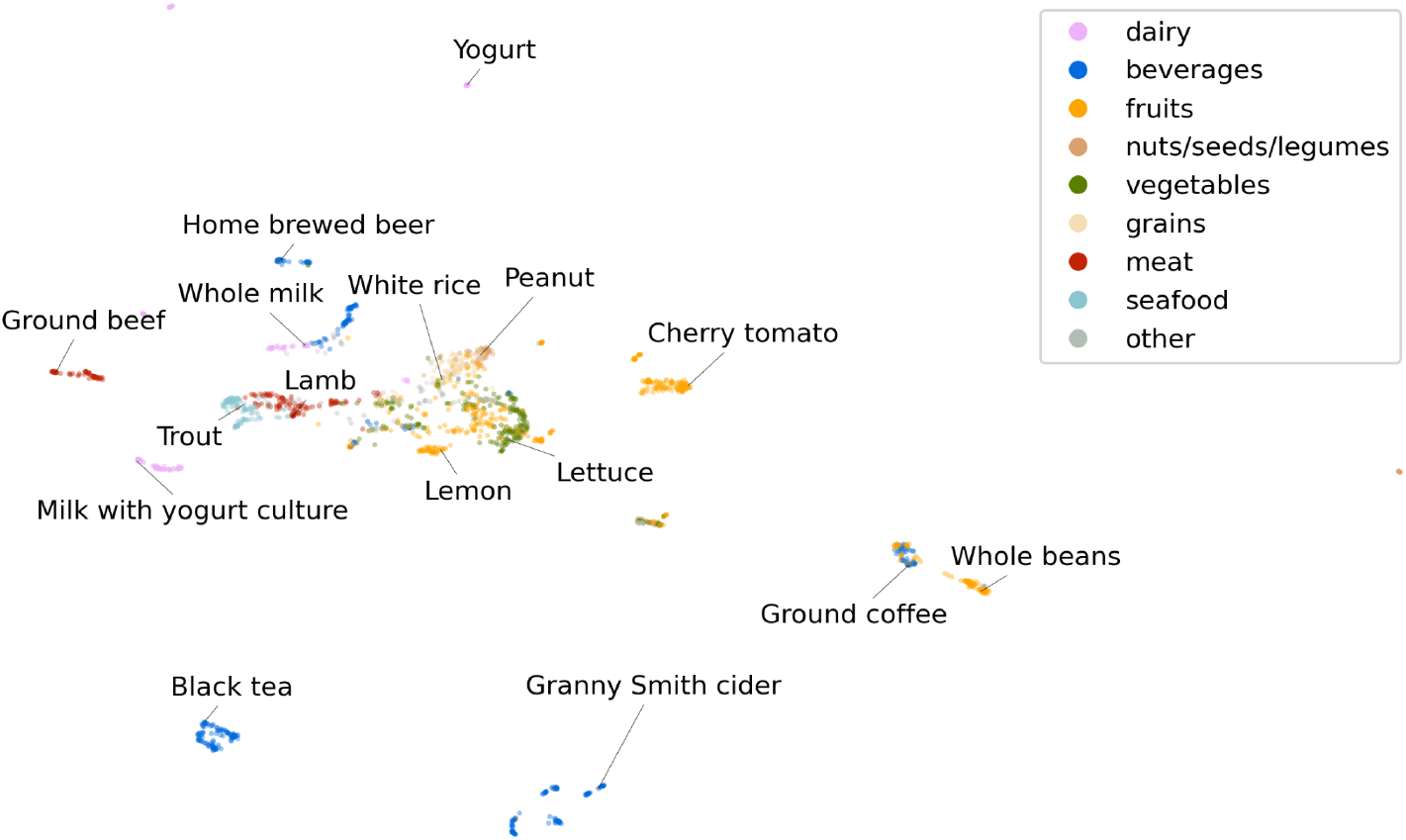
UMAP embedding of untargeted metabolomics data from the Global FoodOmics project based on TreeMS2 spectral similarity. Each point represents one food item, coloured by its assigned food category from curated metadata. Subgroups corresponding to meat, seafood, and dairy are visible, while plant-based items separate into grains/grasses, vegetables/herbs, and fleshy fruits. Selected examples are annotated to illustrate category placement.

In general, major food groups are clustered together, such as seafood, meat, dairy, and vegetables. In addition, animal-based products like seafood and meat are closer together and fruits were clustered close to vegetables (**Figure 6**. and **Supplementary Figure 2**). By annotating the UMAP based on different metadata features, we could illustrate that beverages that contain alcohol like wines and beers are nicely clustered together (**Supplementary Figure 3b**). Whereas the fermented foods cluster separately from their not fermented counterparts, as can be seen for milk and its derivatives like yogurt and cheese (**Supplementary Figure 3c**).

## Discussion

TreeMS2 provides a scalable approach for constructing molecular phylogenies and low-dimensional embeddings directly from raw MS/MS spectra in a broad set of applications ranging from DDA to DIA and from proteomics, to single cell proteomics, and metabolomics. Sequence-based phylogenetics has decades of methodological refinement, but until now there has been no equivalent approach for molecular phenotype data at the scale of modern MS datasets. TreeMS2 bridges this gap, combining scalability to millions of spectra, cross-omics operation, and direct use of raw spectral data, thereby bypassing the limitations imposed by incomplete reference databases and libraries.

Applications across diverse datasets demonstrate both biological relevance and technical robustness. In bacterial and eukaryotic proteomics, TreeMS2 reconstructed established taxonomy down to the genus level, with strong agreement to reference classifications. Deviations from expected placements corresponded to documented sample mix-ups or plausible handling errors, indicating that the method can serve as an unsupervised quality-control check. In the Global FoodOmics dataset, TreeMS2 captured broad biochemical divisions between animal and plant products as well as fine-scale structure within each group from untargeted metabolomics data. In single cell proteomics, TreeMS2 resolved the separation between different cell types and handled noisy, sparse, and variable single-cell DIA data without relying on peptide or protein identifications.

These results establish TreeMS2 as a new type of molecular tool, complementing sequence-based phylogeny approaches by revealing phenotype-level molecular relationships. By capturing functional convergence and divergence across taxa, TreeMS2 can provide insights into adaptation, ecological specialization, and biochemical diversity that are invisible to genomic-only comparisons. More broadly, the method enables large-scale exploration of proteomics and metabolomics datasets for comparative biology, ecology, food science, and biotechnology. The ability to detect sample mix-ups, processing anomalies, and other outliers directly from raw spectra introduces a valuable layer of automated quality control, while unexpected groupings in the resulting trees and embeddings can prompt new hypotheses about shared functional roles or compositional similarities across organisms, environments, or products.

In doing so, TreeMS2 establishes a scalable and domain-agnostic method for quantifying molecular relatedness from MS data. As the volume and diversity of MS data continue to grow, TreeMS2 offers a key capability for linking molecular composition to evolutionary, ecological, and applied contexts.

## Methods

### TreeMS2 workflow

TreeMS2 computes pairwise distances between MS/MS datasets at scale by combining spectral vectorization with approximate nearest-neighbour search (**Figure 7**). Input data consist of one or more MGF (Mascot Generic Format) files and a comma- or tab-separated file mapping each spectrum file to a user-defined group label, allowing multiple runs (e.g., technical replicates or LC–MS/MS fractions) to be aggregated into a single biological unit.

**Figure 7.**
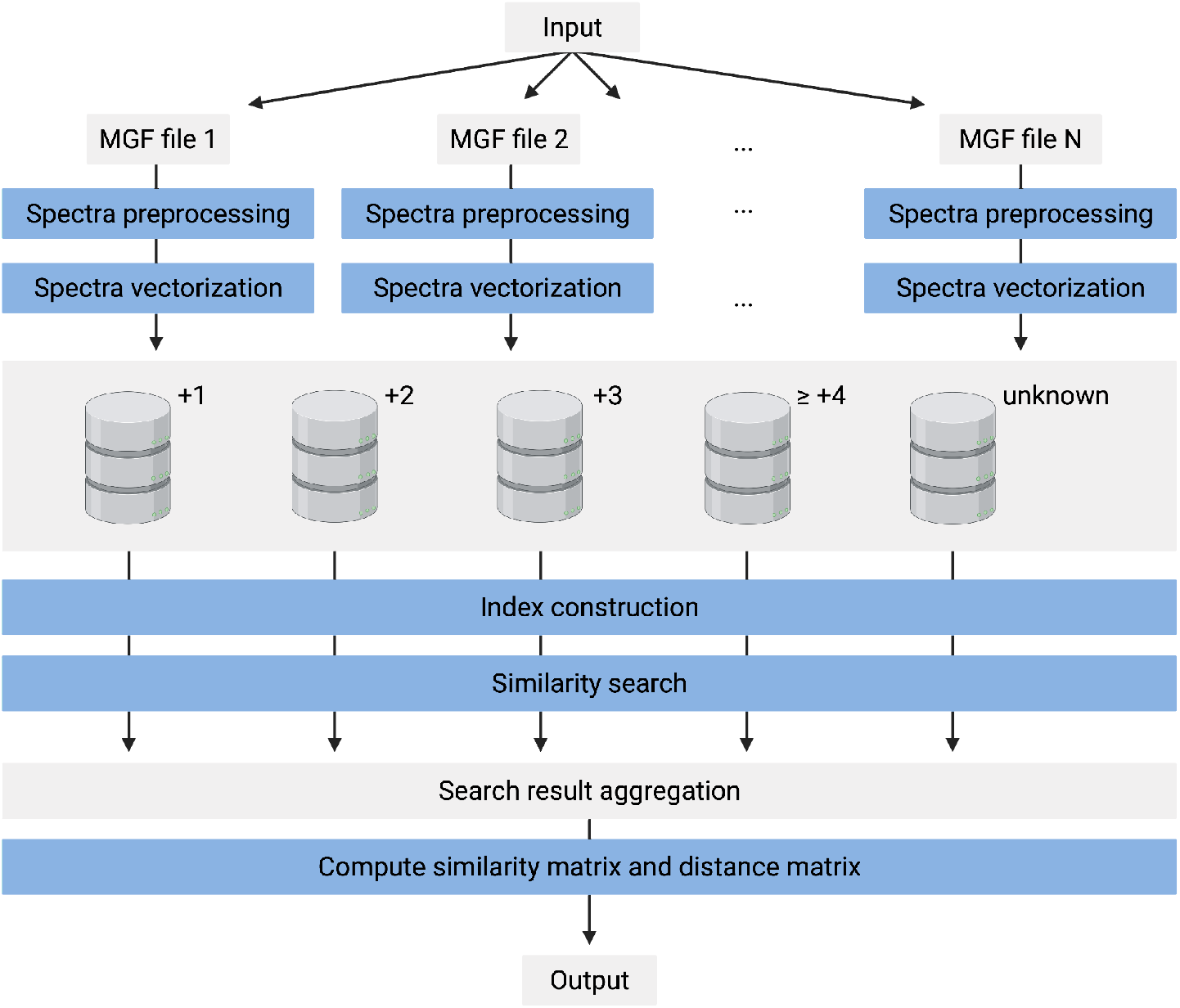
Overview of the TreeMS2 workflow. Input MGF files are processed in parallel, with spectra preprocessed, vectorized, and stored in Lance vector stores partitioned by precursor charge. An approximate nearest neighbor index is built and searched for each store, and results are aggregated across charges to compute a global similarity matrix. This matrix is converted into a distance matrix representing the approximate distances between all user-defined spectral sets.

Spectra are preprocessed to remove noise and ensure consistent representation. Peaks outside a specified *m*/*z* range (default 101–1500 *m*/*z*) or within a precursor-exclusion window (default ±1.5 *m*/*z*) are removed, and low-intensity peaks are filtered relative to the base peak intensity (default ≥1%). Optionally, the top N most intense peaks are retained (default 50). Spectra with fewer than a minimum number of peaks (default 5) or with a total m/z range below a configurable threshold (default 250.0) are discarded. Peak intensities can optionally be scaled using a method of preference (default: square-root) and are subsequently L2-normalized. Each processed spectrum is converted into a high-dimensional sparse vector by binning fragment *m*/*z* values at a specified tolerance (default 0.05 *m*/*z*). Intensities within the same bin are summed. To reduce dimensionality while approximately preserving cosine similarity, TreeMS2 applies sparse random projection [27] to generate dense vectors of configurable size (default 400) that are subsequently L2-normalized.

For proteomics data, the default settings as described above were used. For metabolomics data, where acquisition characteristics differ, preprocessing was adapted by lowering the minimum m/z to 50 and the minimum required m/z range to 100.

Vectors are stored in Lance [28] vector stores, partitioned by precursor charge (1, 2, 3, ≥4, and undefined) to restrict comparisons to spectra with similar fragmentation patterns. Each store is indexed separately using Faiss [29], with the index type selected dynamically based on dataset size. For small stores (<10,000 vectors), a flat exhaustive search index is used. For larger stores, TreeMS2 constructs inverted file (IVF) indexes [30], where the number of clusters is proportional to the square root of the number of vectors, ensuring a balance between index resolution and centroid robustness. Centroids are computed via k-means clustering on a representative sample (≥39 vectors per cluster). If a graphics processing unit (GPU) is available, it can be used to accelerate the computation. For very large indexes (≥1 million vectors), hierarchical navigable small world (HNSW) graphs [31] are used as coarse quantizers to accelerate centroid search, avoiding the need to compare against all centroids directly.

TreeMS2 monitors available system memory and estimates a per-vector memory budget from the total memory, the current usage of the process, and a reserved margin for the operating system. Depending on this budget, vectors may be stored uncompressed as 32-bit floats, stored in half-precision (16-bit floats) for moderate compression, quantized to 8 bits per component for further reduction, or compressed via product quantization [32], which divides vectors into subvectors and quantizes them independently. This adaptive approach ensures that large datasets can be indexed in systems with limited memory, while maintaining search accuracy as far as possible.

For each spectrum, the index is queried for a user-defined number of nearest neighbours (default 1024) across a set number of clusters (default 128). Only neighbours above a similarity threshold (default 0.8) and within an optional precursor *m*/*z* window (default 2.05 *m*/*z*) are retained. Search results are aggregated across batches and precursor-charge partitions to count, for each input group pair, the fraction of spectra in one with at least one match in the other.

Using the aggregated matrix from the previous step, TreeMS2 computes the global similarity and global distance between all sets of spectra, as explained in section 1.2.2. The result is a symmetric distance matrix, where each entry represents the (approximate) distance between two sets. Since the distance between a set and itself is always zero and the matrix is symmetric, it can be represented as a strictly lower triangular matrix. TreeMS2 outputs a distance matrix in the MEGA (.meg) file format, which is a human-readable format used by the MEGA software suite (Molecular Evolutionary Genetics Analysis) [33]. This file can be opened in MEGA (version 11) to construct a rooted tree based on the distance matrix, using the UPGMA algorithm.

### Comparison between TreeMS2 and compareMS2

TreeMS2 was compared to the current state-of-the-art compareMS2 (version 2.1.0) [8] on 13 files selected at random from the Pseudomonas dataset and 7 files from the Bacillus dataset, both available on MassIVE MSV000096603 (**Supplementary Table 4**). To assess the scalability of the tools, the number of spectra to be compared were gradually increased by gradually adding more files. For detailed information on data acquisition, please refer to the original publication by Abele et al. [17]. In brief, bacteria were lysed and 50 μg proteins were reduced, alkylated, and digested with trypsin. Around 500 ng of peptides were spiked with 50 fmol retention time standardization peptides. The peptides were analyzed on an UltiMate 3000 RSLCnano (Thermo Fisher Scientific) coupled to an Orbitrap Fusion Lumos Tripbrid mass spectrometer (Thermo Fisher Scientific). The mass spectrometer was operated in DDA and positive ionization mode.

### Use-case 1: Bacterial proteomics dataset

For this use-case the 38SPD diversity dataset from Abele et al. [17] was selected. This dataset contains the proteomic data of 318 strains from 303 species measured with an LC-gradient of 30 minutes. This dataset contains 340 files and is available on MassIVE: MSV000096603.

For detailed information on data acquisition, please refer to the original publication by Abele et al. [17]. All cultures for the diversity datasets were obtained from strains stored in the Weihenstephan Microbial Strain Collection (https://ccinfo.wdcm.org/details?regnum=1163). Bacteria were lysed and 20μg proteins were reduced, alkylated, and digested with trypsin. 10μg of peptides were analyzed on a Vanquish Neo UHPLC (microflow configuration; Thermo Fisher Scientific) coupled to an Orbitrap Exploris 480 mass spectrometer (Thermo Fisher Scientific). The mass spectrometer was operated in DDA and positive ionization mode. Sequence database searching was performed with MSFragger (version 4.1) with default parameters against species-specific fasta files.

### Calculate taxonomic relatedness

To assess taxonomic relatedness to a sequence-based phylogenetic tree. Our tree obtained by analyzing the 38SPD dataset [17] was compared to the GTDB dataset (https://lpsn.dsmz.de/) by using a Mantel test with Spearman correlation. 295 of the 302 species present in the 38SPD dataset were present in the GTDB dataset (https://lpsn.dsmz.de/). The 7 missing species were not included in the taxonomic relatedness analysis. The species mapping across 38SPD and GTDB can be found in **Supplementary Table 5**. To obtain values per species rather than per sample, the similarity matrix was aggregated to the species level by averaging the pairwise distances among samples belonging to the same species.

### Food-derived bacterial isolates classification

To assess the ability of TreeMS2 to classify observed bacteria in food, we analyzed the food isolate dataset with the 38SPD dataset. The food dataset contains 60 dairy product isolates measured with an LC-gradient of 10 minutes. For detailed information on data acquisition, please refer to the original publication by Abele et al. [17]. In brief, the isolates from dairy products were sampled directly from agar plates. The bacteria were lysed and 20μg proteins were reduced, alkylated, and digested with trypsin. 10μg of peptides were analyzed on a Vanquish Neo UHPLC (microflow configuration; Thermo Fisher Scientific) coupled to an Orbitrap Exploris 480 mass spectrometer (Thermo Fisher Scientific). The mass spectrometer was operated in DDA and positive ionization mode. The data are available on MassIVE: MSV000096603.

Of the 60 food isolates, 7 were excluded: 2 were QC runs, 3 had no 16S rRNA results, and 2 had incomplete results. Of the remaining 53 food isolates, 46 had their genus represented in the 38SPD dataset, of which 32 even had the species represented in the 38SPD dataset. Genus classification was obtained through a nearest neighbor classifier, looking at the 5 closest neighbors and calculating a weighted majority vote based on the distance. Once the genus was established, the closest neighbour that matches with the predicted genus is reported as the matched species.

### Use-case 2: kingdom of life

To evaluate TreeMS2 on larger and more complex proteomes, we analyzed data from a “kingdom of life” proteomics study [21], containing 14 viruses, 19 archaea, 49 bacteria, and 32 eukaryotes, a total of 114 species. For detailed information on data acquisition, please refer to the original publication by Müller et al. [21]. In brief, the in-StageTip (iST) protocol [34] was used for automated and reproducible sample preparation. To reduce the complexity in the measurements, eukaryotic peptide mixtures were separated into eight fractions. All samples were measured on an EASY-nLC 1200 ultrahigh-pressure system (Thermo Fisher Scientific) coupled to a Q Exactive HFX Orbitrap instrument (Thermo Fisher Scientific) with a nano-electrospray ion source (Thermo Fisher Scientific). The mass spectrometer was operated in DDA and positive ionization mode. The data are available on PRIDE: PXD014877.

In total, 381 RAW files were available. However, 52 files could not be converted to MGF format using ThermoRawFileParser (version 1.4.5) [35], resulting in 329 successfully converted MGF files. Additionally, not all files could be reliably mapped to their corresponding species. Consequently, instead of 14 viruses, 19 archaea, 48 bacteria, and 33 eukaryotes, only 79 species remained (11 viruses, 3 archaea, 32 bacteria, and 33 eukaryotes). A full overview of the included species and RAW files can be found in Supplementary Table 2.

*E. coli* was the food source on the agar plate *D. discoideum* was growing on (Carolina item #155996). To investigate whether *E. coli* was sampled together with *Dictyostelium discoideum*, a database search was performed on eight *D. discoideum* fractions using MSFragger (version 4.1) with default parameters. A combined fasta database containing *E. coli* and *D. discoideum* protein sequences was used.

The TreeMS2 tree is compared to the NCBI taxonomy tree. To incorporate some basic distance information to the NCBI taxonomy tree, a distance was calculated based on how many taxonomic rank nodes agree between two neighbouring species.

### Use-case 3: Single Cell Proteomics data

To illustrate TreeMS2 on a DIA application, the tool was applied on a SCP dataset. For detailed information on data acquisition, please refer to the original publication by Ye et al. [23]. In brief, hi12iPSCs were cultured and induced with EB induction. Samples were lysed and digested within the proteoCHIP EVO 96 inside the cellenONE (Cellenion SASU). The cell isolation process uses the precision of the cellenONE module to sort individual cells, which are selected based on morphological criteria (diameter range of 22–30 µm and a maximum elongation factor of 1.6), into each well. An Evosep One chromatography system (Evosep Biosystems) was coupled to an Orbitrap Astral mass spectrometer (Thermo Fisher Scientific). The mass spectrometer was operated in DIA and positive ionization mode. The data were downloaded from the PRIDE repository with identifiers PXD014877 and PXD019483.

Multidimensional scaling (MDS) was applied to obtain a low-dimensional embedding of the samples. The distance matrix generated by TreeMS2 was used directly as a dissimilarity matrix. Classical metric MDS was performed using the scikit-learn implementation (version 1.4.1.post1) and the first two dimensions were retained for visualization. The cluster annotation from the original paper by Ye et al. [23] was used.

### Use-case 4: Global FoodOmics data

TreeMS2 was also applied on a large untargeted metabolomics dataset. As certain parameters for metabolomics data acquisition differ from proteomics data acquisition, the spectrum preprocessing in TreeMS2 was adapted for the metabolomics context. Specifically, the minimum m/z was set to 50 rather than 101 in the proteomics context, and the minimum m/z range was set to 100 rather than 250.

For detailed information on data acquisition, please refer to the original publication by Gauglitz et al. [25]. In brief, metabolites were extracted through ethanol extraction from 3,579 food and beverage samples. Food extracts were analyzed using an UltiMate 3000 ultra-high-performance liquid chromatography system (Thermo Scientific) equipped with a reverse phase C18 column coupled to a Maxis Q-TOF Impact II mass spectrometer (Bruker Daltonics) equipped with an electrospray ionization source. The data were downloaded from the MassIVE repository with identifier MSV000084900.

All foods are curated and include a six-level food ontology, as well as information for fermentation or organic status, land or aquatic origin, and country of origin. For this case-study, we selected all simple foods. Resulting in a diverse set of foods from different categories such as dairy, fruit, meat, seafood, and vegetables.

Uniform Manifold Approximation and Projection (UMAP) was applied to obtain a low-dimensional embedding of the samples. The precomputed pairwise distance matrix was used directly as a dissimilarity matrix. UMAP was performed using the umap-learn implementation (version 0.5.11) with the metric set to “precomputed” and the number of nearest neighbors set to 50. The first two UMAP dimensions were retained for visualization. Sample annotations from the associated metadata table were used to color the embedding according to the specified category labels.

### Software availability

The TreeMS2 source code is available on https://github.com/bittremieuxlab/TreeMS2.

