## Supplementary Figures and Tables for "Scalable mass-spectrometry-based molecular phylogeny with TreeMS2"

**Supplementary Figure 1. Bacterial proteome full tree.**

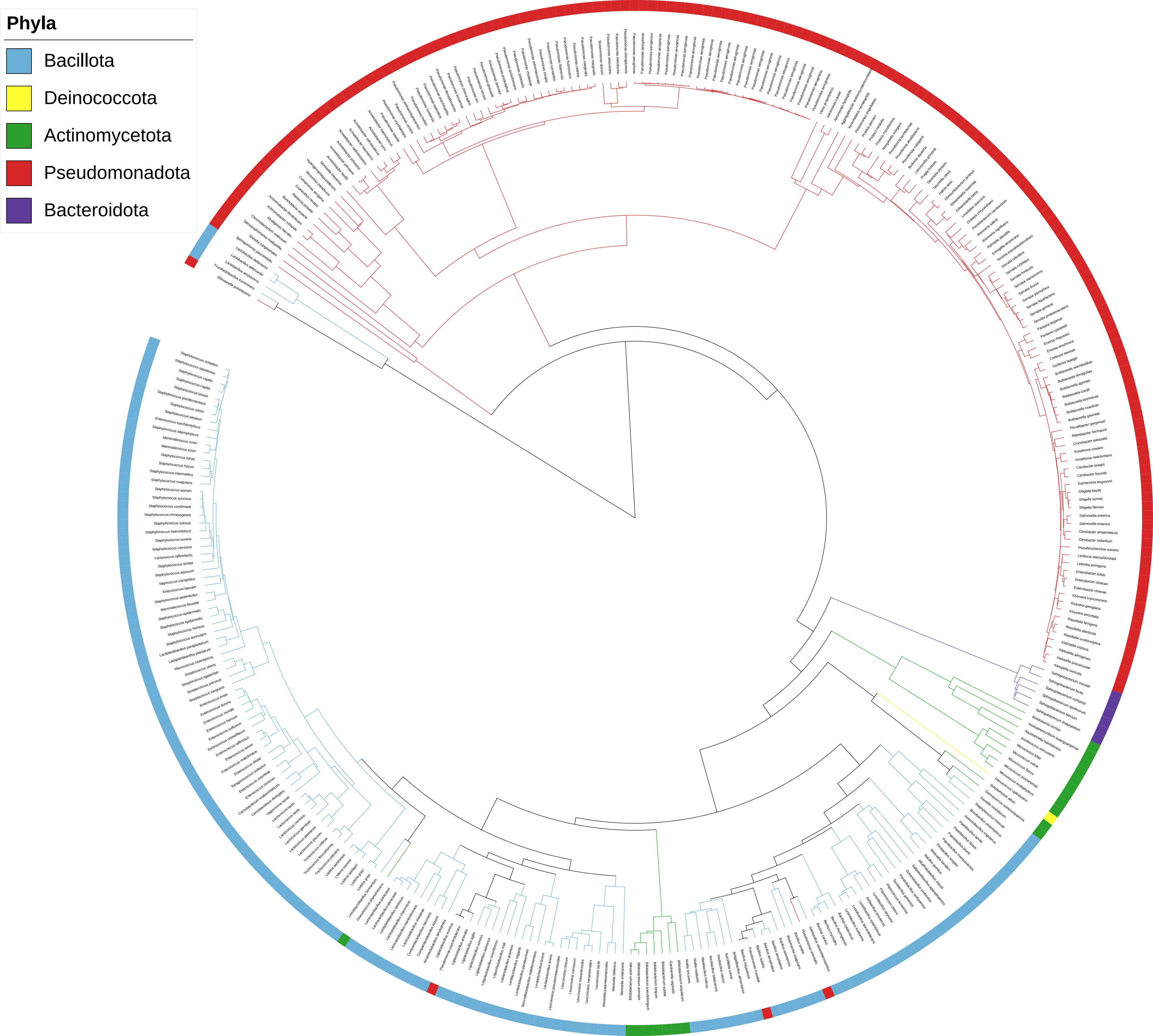
Phyla
Bacillota
Deinococcota
Actinomycetota
Pseudomonadota
Bacteroidota

**Supplementary Figure 2. FoodOmics full tree.**

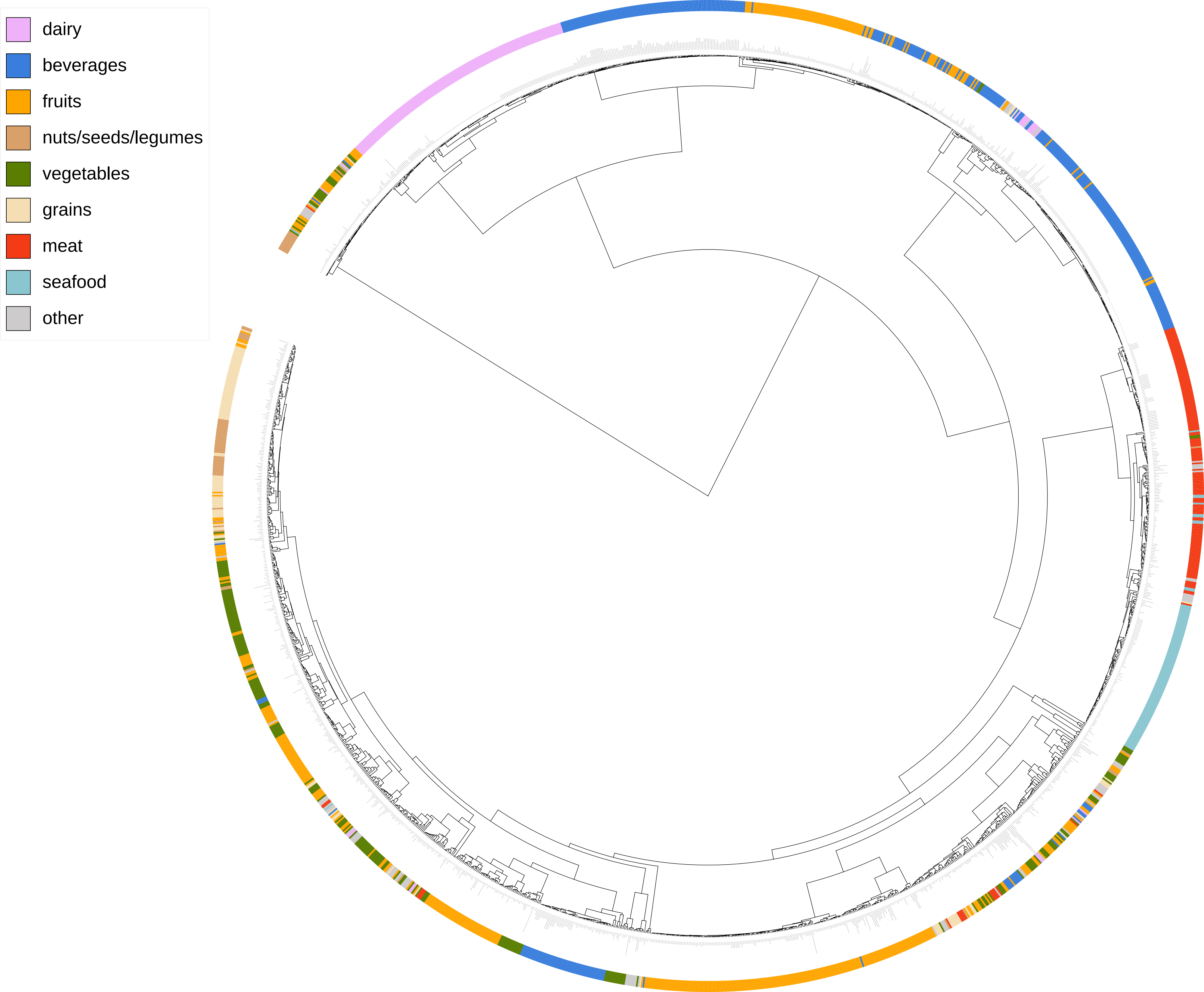
dairy
beverages
fruits
nuts/seeds/legumes
vegetables
grains
meat
seafood
other

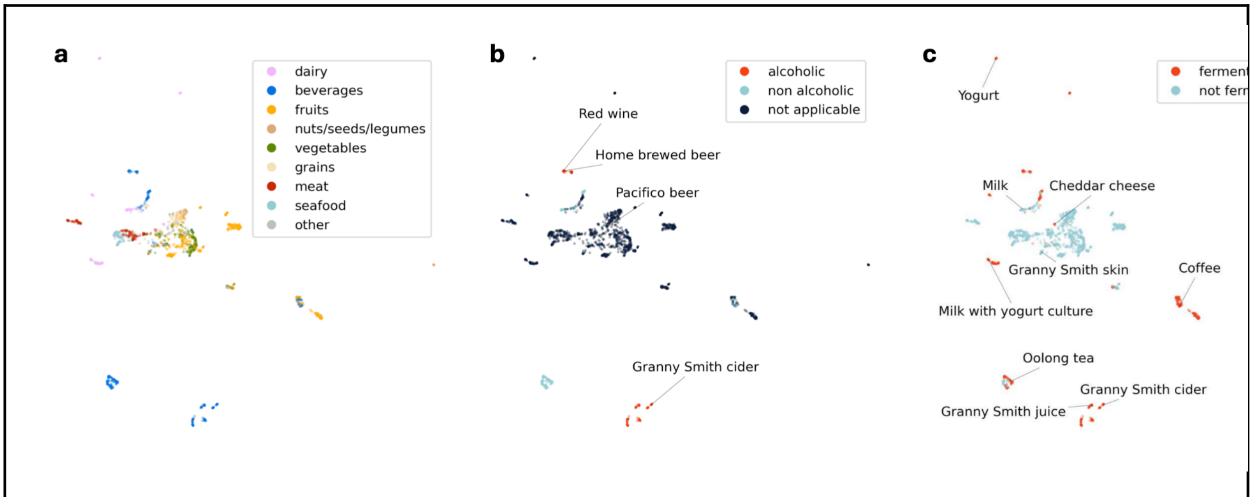


**Supplementary Figure 3. UMAP embedding of untargeted metabolomics data from the Global FoodOmics project based on TreeMS2 spectral similarity.** Each point represents one food item. The foods are coloured based on curated metadata by its assigned food category (**a**), their alcohol content (**b**), or whether they are fermented (**c**).

| Dataset | Num_peak_files | Num_spectra | Num_retained | Num_removed | Runtime (s) |
|---|---|---|---|---|---|
| 38SPD | 340 | 13808820 | 13663202 | 145618 | 12016 |
| 38SPD dairy isolates | 400 | 14691952 | 14533344 | 158608 | 17266 |
| foodomics | 1859 | 4531245 | 1597256 | 2933989 | 5037 |
| iPSC_EBs | 104 | 21146548 | 20876393 | 270155 | 12091 |
| kingdom of life | 326 | 56122054 | 55730897 | 391157 | 45127 |

**Supplementary Table 1. Runtime information.** This table contains the number of processed spectra and runtime for each dataset on which TreeMS2 was applied.

**Supplementary Table 2** (sup-table-2-kingdom-of-life-included-species-overview.tsv): This table contains an overview of which species were included from the kingdom of life dataset.

**Supplementary Table 3** (sup-table-3-kingdom_of_life_tanglegram-neighbour_comparison_summary.tsv): This table contains how many neighbours match between the TreeMS2 tree and the NCBI tree.

**Supplementary Table 4** (sup-table-4-compareMS2-comparison-files-used.tsv): List of file names used for TreeMS2 - CompareMS2 comparison.

| Files | Dataset |
|---|---|
| BBM_334_P110_25_DDA_011.raw | Pseudomonas |
| BBM_338_P110_28_DDA_003.raw | Bacillus |
| BBM_338_P110_28_DDA_004.raw | Bacillus |
| BBM_334_P110_25_DDA_010.raw | Pseudomonas |
| BBM_338_P110_28_DDA_010.raw | Bacillus |
| BBM_334_P110_25_DDA_21_R2.raw | Pseudomonas |
| BBM_338_P110_28_DDA_014.raw | Bacillus |
| BBM_334_P110_25_DDA_015.raw | Pseudomonas |
| BBM_334_P110_25_DDA_005_R2.raw | Pseudomonas |
| BBM_334_P110_25_DDA_020.raw | Pseudomonas |
| BBM_334_P110_25_DDA_018.raw | Pseudomonas |
| BBM_334_P110_25_DDA_012.raw | Pseudomonas |
| BBM_334_P110_25_DDA_028.raw | Pseudomonas |
| BBM_334_P110_25_DDA_006.raw | Pseudomonas |
| BBM_334_P110_25_DDA_019.raw | Pseudomonas |
| BBM_338_P110_28_DDA_009.raw | Bacillus |
| BBM_334_P110_25_DDA_001_R2.raw | Pseudomonas |
| BBM_338_P110_28_DDA_016.raw | Bacillus |
| BBM_334_P110_25_DDA_016.raw | Pseudomonas |
| BBM_338_P110_28_DDA_012.raw | Bacillus |

**Supplementary Table 4. List of file names used for TreeMS2 - CompareMS2 comparison.** This table contains the files used to compare the runtime efficiency between TreeMS2 and CompareMS2. Files were added in the same order as specified in the table.
